# An intra-family conserved high-order RNA structure within the M ORF is important for arterivirus subgenomic RNA accumulation and infectious virus production

**DOI:** 10.1101/2024.05.30.596123

**Authors:** Pengcheng Shang, Yanhua Li, Chi Chen, Ian Brierley, Andrew E. Firth, Ying Fang

**Author notes:** These authors contribute equally to the study (Pengcheng Shang, Yanhua Li). Children’s Hospital of Pittsburgh, Pittsburgh, USA.

## Abstract

Synthesis of subgenomic RNAs is a strategy commonly used by polycistronic positive sense single-stranded RNA viruses to express 3′-proximal genes. Members of the order of *Nidovirales*, including coronaviruses and arteriviruses, use a unique discontinuous transcription strategy to synthesize subgenomic RNAs. In this study, *in silico* synonymous site conservation analysis and RNA structure folding predicted the existence of intra-family conserved high-order RNA structure within the M ORF of arteriviral genomes, which was further determined to be important for the transcription/accumulation of subgenomic RNAs and production of infectious viral particles. Mutations disrupting the stability of the RNA structures significantly decreased the accumulation of multiple subgenomic RNAs. In contrast, the impact of mutagenesis on full-length genomic RNA accumulation was limited. The degree to which wild-type levels of subgenomic RNA accumulation were maintained was found to correlate with the efficiency of infectious virus production. Moreover, the thermo-stability of stems within the high-order RNA structure is also well correlated with viral replication capacity and the maintenance of subgenomic RNA accumulation. This study is the first to report an intra-*Arteriviridae* conserved high-order RNA structure that is located in a protein-coding region and functions as an important *cis*-acting element to control the accumulation/transcription of arteriviral subgenomic RNAs. This work suggests a complex regulation mechanism between genome replication and discontinuous transcription in nidoviruses.

**IMPORTANCE:** Arteriviruses are a group of RNA viruses that infect different animal species. They can cause diseases associated with respiratory/reproductive syndromes, abortion, or haemorrhagic fever. Among arteriviruses, porcine reproductive and respiratory syndrome virus (PRRSV) and equine arteritis virus (EAV) are economically important veterinary pathogens. The challenge in control of arterivirus infection reflects our limited knowledge of viral biology. In this study, we conducted a comprehensive bioinformatical analysis of arteriviral genomes and discovered intra-family conserved regions in the M ORF with a high-order RNA structure. The thermo-stability of the RNA structure influences sgRNA transcription/accumulation and correlates with the level of infectious virus production. Our studies provide a new insight on arterivirus replication mechanism, which may have implications in developing disease control and prevention strategies.

## INTRODUCTION

For polycistronic positive sense (+) single-stranded (ss) RNA viruses, the synthesis of subgenomic RNAs (sgRNAs) is a common strategy to allow ribosomal access to genes that are encoded in 3′ regions of the genomic RNA (1, 2). In addition, as independently transcribed products, sgRNAs have the potential to be regulated during virus infection in a spatial, temporal and/or quantitative manner (1, 2).

In members of the order *Nidovirales*, the synthesis of sgRNAs is essential for the expression of structural and accessory proteins encoded in 3′-proximal ORFs (3, 4). Nidoviral subgenomic transcripts are 3′ co-terminal with the full-length genomic RNA (gRNA). For arteriviruses and coronaviruses, the 5′ terminal sequence of sgRNAs is known as the leader sequence and is identical to the 5′ end of the gRNA (3, 4). Two mechanisms have been proposed to explain the synthesis of these nidoviral sgRNAs (3). In the leader-primed transcription model, plus-strand (+) sgRNAs are discontinuously transcribed from full-length minus-strand (−) gRNAs. In contrast, in the more widely accepted discontinuous minus-strand RNA synthesis model, (+) sgRNAs are synthesized from template (−) sgRNAs, which are transcribed discontinuously from full-length (+) gRNAs and/or longer (+) sgRNAs (3–5). At the end of the leader sequence of each (+) sgRNA, there is a short conserved AU-rich transcription regulatory sequence (TRS), which is believed to play important roles in mediating discontinuous transcription. The TRS present immediately downstream of the leader sequence within the full-length (+) gRNA is named the leader TRS (L-TRS), whereas the TRSs found upstream of 3′-proximal ORFs are called body TRSs (B-TRSs). The discontinuous transcription of (−) sgRNAs is guided by the base pairing interaction between a newly synthesized antisense (−) B-TRS with a sense (+) L-TRS. Every B-TRS would serve as an attenuation signal to polymerase processivity, with (−) RNA synthesis either able to continue on the (+) gRNA template, or else the nascent (−) RNA disassociates from the (+) gRNA template at the B-TRS and reanneals to a (+) gRNA template at the L-TRS, where polymerase activity resumes to synthesize the antisense leader sequence templated by the 5′-proximal region of the (+) gRNA. During replication, the replication and transcription complex (RTC) reads through all B-TRSs without interruption to synthesize a full-length (−) gRNA (3–5). The (+) gRNA to (−) gRNA to (+) gRNA cycle is traditionally designated as replication, whereas the (+) gRNA to (−) sgRNAs to (+) sgRNAs step is referred to as transcription (3, 4).

Like other members of the order *Nidovirales*, synthesis of arterivirus gRNAs and discontinuous transcription of sgRNAs has been demonstrated to be precisely regulated by both *cis*-acting and *trans*-acting factors (3–7). The identity of the primary sequence of the B-TRS and the stability of the B-TRS : L-TRS duplex have been found to correlate with the relative production levels of different sgRNAs (3–7). In the equine arteritis virus (EAV), the L-TRS is presented within the loop of an RNA stem-loop structure, which was characterized to facilitate the (−) B-TRS : (+) L-TRS duplex formation. Its important role in discontinuous transcription has been elucidated in site-directed mutagenesis studies (8–13). In addition to *cis*-acting RNA signals, specific replicases were determined to function as *trans*-acting factors to influence the activity of the RNA-dependent RNA polymerase (RdRp) in RNA transcription, as demonstrated in EAV studies (4, 14, 15). Subgenomic RNA transcription was nearly abolished by one single mutation in EAV nsp10 (helicase), while full-length genome synthesis and nonstructural polyprotein processing remain unaltered (15, 16). Additionally, mutagenesis studies revealed that the EAV nsp1 protein is a prominent *trans*-acting factor to ensure the equilibrium of sgRNA transcription and genome replication, and maintaining the production ratios of different sgRNA species (14).

So far, potential *cis*-acting elements within protein-coding regions, which are able to modulate arterivirus subgenomic RNA transcription, are largely unknown. In this study, an in-depth bioinformatics analysis revealed a specific region located within the M ORF of arteriviral genomes which has a significantly reduced rate of substitutions at synonymous sites, indicative of an overlapping functional element. This region was predicted to form a potential high-order RNA structure, including two short stem-loops (SL1 and SL2) and an extended stem-loop (extSL). One member of the *Arteriviridae* family, porcine reproductive and respiratory syndrome virus (PRRSV), was used to investigate the function of this RNA structure in the virus life cycle. The results demonstrate that this conserved high-order RNA structure in the M ORF plays an important role in controlling the accumulation/transcription of arteriviral subgenomic RNAs.

## MATERIALS AND METHODS

### Synonymous site conservation analysis

Synonymous site conservation was analyzed with synplot2 (17). First, we used tblastn (18) to identify PRRSV-1 and PRRSV-2 sequences in the NCBI nr/nt database (accessed between 17th and 20th February 2023; taxa restricted to *Riboviria* i.e. txid2559587). We used the translated concatenated coding regions of NCBI RefSeqs NC_043487 and NC_001961 (for PRRSV-1 and PRRSV-2, respectively) as the tblastn query sequences; in regions of gene overlap, the reading frame selected for translation was the reading frame of the longer of the overlapping ORFs. Sequences were selected for alignment if they had > 95% coverage and > 75% amino acid identity to the RefSeq concatenated coding regions. There were no sequences in common between the sequences matched to the PRRSV-1 RefSeq and the sequences matched to the PRRSV-2 RefSeq. To reduce computation, 200 of the retrieved sequences were then selected at random for each of PRRSV-1 and PRRSV-2. Full-genome codon-respecting alignments were generated as described in previous work (17). In brief, each individual genome sequence was aligned to the reference sequence (NC_043487 or NC_001961) using code2aln version 1.237 (19). Genomes were then mapped to the reference sequence coordinates by removing alignment positions that contained a gap character in the reference sequence, and these pairwise alignments were combined to give the multiple sequence alignment. To assess conservation at synonymous sites, coding regions were extracted from the alignment (with codons selected from the longer ORF in each overlap region), concatenated in-frame, and the alignment analysed with synplot2 using a 9-codon sliding window. Conservation statistics were then mapped back to reference genome coordinates for plotting (Fig. 1).

**Figure 1.**
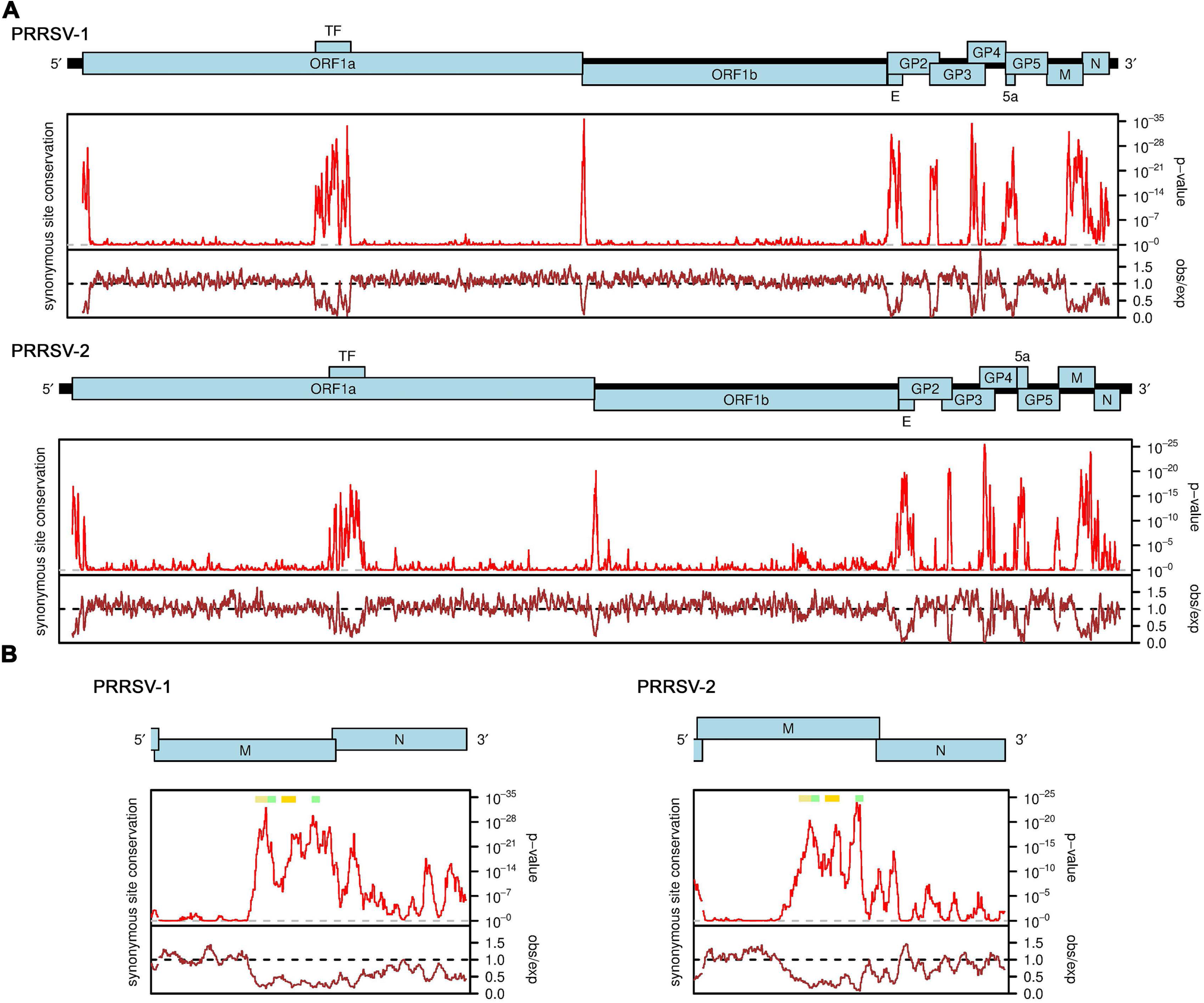
Synonymous site conservation analysis of PRRSV sequences. **(A)** Full genome analyses for PRRSV-1 (upper) and PRRSV-2 (lower) sequence alignments. The red line shows the probability that the observed conservation could occur under a null model of neutral evolution at synonymous sites, whereas the brown line depicts the ratio of the observed number of substitutions to the number expected under the null model. Note that similar plots for PRRSV-2 (albeit with different input sequences) have been presented previously in Firth (2014) and Cook et al. (2022). **(B)** Expanded views of the M and N ORF region. Positions of predicted RNA structures are indicated with colored bars: yellow – stem-loop 1, SL1; orange – stem-loop 2, SL2; green – 5′ and 3′ fragments of the extended stem-loop, extSL.

### RNA structure prediction

RNA structures were predicted using a combination of manual inspection, RNAfold version 2.4.1 from the ViennaRNA package (20) and RNAStructure (21). RNAz was used via the ViennaRNA online server (http://rna.tbi.univie.ac.at) on 29 Dec 2023 using default parameters except that the ’Target mean pairwise identity’ was changed from 80% to 65%. The input was a full genome alignment constructed using code2aln version 1.237 (19) of the sequences NC_001961, NC_028963, NC_040535, NC_043487, NC_048210 and NC_075972 (since RNAz takes a maximum of six sequences, the seventh sequence NC_032987 was omitted). In Fig. 5, energies for wild-type and mutant SL1 and SL2 were calculated with RNAfold whereas energies for wild-type and mutant extSL were calculated with RNAduplex (version 2.4.1, ViennaRNA package), using the sequence regions shown as input.

### Phylogenetic analysis

We downloaded all arterivirus sequences (txid76803) of length > 10000 nt from the NCBI nr/nt database on 12 Dec 2023. We discarded sequence records containing the "KEYWORDS UNVERIFIED" flag (often low-quality sequences) or "PAT" or "SYN" in the "LOCUS" line (patent and synthetic sequences), and sequences containing > 20 ambiguous nucleotide codes. Next, we extracted the ORF1ab regions using the criteria: an AUG-to-stop-codon ORF with length > 4000 nt and starting not later than nt 2000 of the genome as ORF1a; the last U_UUA_AAC upstream of the ORF1a stop codon as the ribosomal frameshift site; and the ORF measuring from the C of U_UUA_AAC to the next in-frame stop codon, with length >3000 nt as ORF1b. Sequences that did not contain ORFs 1a and 1b meeting these thresholds were discarded, except for the partial sequence MT085119 (in which ORF1a is 5’-truncated to 2880 nt due to missing 5’ sequence data) which was retained due to the absence of any closely related more-complete sequence. Following the phylogenetic trees presented in previous studies (22–24), we excluded the divergent equine arteritis virus (genus *Alphaarterivirus*) and wobbly possum disease virus (genus *Kappaarterivirus*) clades, and a couple of more divergent sequences (isolated from reptiles). We clustered the ORF1ab amino acid sequences using BlastClust (18) with parameters -L 0.90 -b F -S 90 (i.e. 90% identity and 90% coverage thresholds) resulting in 93 clusters. The singleton clusters containing OP168793.1 and OR711915.1 were removed since the ORF1ab polyproteins of these two sequences contain long stretches which appear to align out-of-frame to the ORF1ab region of other PRRSV viruses. A single representative sequence was chosen from each of the remaining 91 clusters – either an NCBI RefSeq if present, else the sequence closest to the centroid (minimum summed pairwise amino acid distances from sequence *i* to all other sequences *j* within the cluster). The ORF1ab amino acid sequences of the representative sequences were aligned with MUSCLE (25) and phylogenetic trees were estimated with PhyML version 3.1 (26) with default parameters for amino acid sequences. An initial phylogenetic tree was produced using wobbly possum disease virus as an outgroup to check there were no other sequences more divergent from PRRSV than Olivier’s shrew virus (genus *Muarterivirus*). Subsequently, a revised phylogenetic tree was produced without wobbly possum disease virus, this time rooted using Olivier’s shrew virus as the outgroup. The tree was then visualised with FigTree (http://tree.bio.ed.ac.uk/software/figtree/).

### Sequences of SL1, SL2 and extSL

At the 90% ORF1ab amino acid identity threshold, BlastClust produced 23 PRRSV-1 and 22 PRRSV-2 clusters and, when combined, these clusters contained 461 PRRSV-1 and 1518 PRRSV-2 genome sequences. For these sequences, we extracted the region from the ORF1b stop codon to the end of the genome sequence, aligned the nucleotide sequences within each of the PRRSV-1 and PRRSV-2 clades using MUSCLE (25), manually checked that the SL1, SL2 and extSL sequences were correctly aligned, and then extracted the regions corresponding to the SL1, SL2 and extSL duplexes from all sequences in each alignment (Fig. 3). For Figs 4, S2, and S3, we extracted the M ORF region from the 91 representative sequences and, within each of two clades – (i) LDV, PRRSV-1, PRRSV-2 and relatives, and (ii) subfamily *Simartivirinae* – we aligned M ORF amino acid sequences with MUSCLE and used the amino acid alignment to guide a codon alignment. SL1, SL2 and extSL predictions were annotated manually.

### Cells

BHK-21 and MARC-145 cells were maintained at 37 °C with 5% CO_2_ in minimum essential medium (MEM) (Gibco, Carlsbad, CA) supplemented with 10% fetal bovine serum (Sigma-Aldrich, St. Louis, MO), 100 U/ml of penicillin plus 100 µg/ml of streptomycin (Gibco, Carlsbad, CA), and 0.25 µg/ml amphotericin B (Fungizone; Gibco, Carlsbad, CA). Infected MARC-145 cells were maintained in 2% horse serum (HyClone, Logan, UT) at 37 °C with 5% CO2.

### Construction and production of PRRSV mutants

PRRSV-1 strain SD01-08 infectious clone-pT7-SD01-08 was constructed in our previous study (27). Site mutagenesis on the M ORF RNA structure was performed as previously described (28, 29). Briefly, the DNA of full-length infectious clone was digested at unique restriction sites-*Xba*I and *Not*I. Two PCR reactions (PCR1 and PCR2) were performed to amplify the genomic region between *Xba*I and *Not*I. The overlapping part of the 3′ end of the PCR1 product and the 5′ end of the PCR2 product contains the mutation sites, which were introduced by primers synthesized from Integrated DNA Technologies (Coralville, IA). The 5′ end of the PCR1 product and the 3′ end of the PCR2 product were overlapped with the infectious clone backbone. These three DNA fragments were further assembled using the NEBuilder HiFi DNA Assembly cloning kit (New England BioLabs, Ipswich, MA).

To generate full-length viral gRNAs for transfection into cells, *in vitro* RNA transcription was performed. The pT7-SD01-08 plasmid DNA was linearized by *Xba*I, which is located immediately downstream of the HDV ribozyme. Digested DNA was extracted using phenol extraction and ethanol precipitation method. *In vitro* RNA transcription was performed in 50 µl reactions: 5 µl RNAPol Reaction Buffer (New England BioLabs, Ipswich, MA), 2 µl T7 RNA Polymerase (New England BioLabs, Ipswich, MA), 5 µl ATP (10 μmol), 5 µl CTP (10 μmol), 5 µl UTP (10 μmol), 2.5 µl GTP (10 μmol) (New England BioLabs, Ipswich, MA), 7.5 µl m7G(5′)ppp(5′)G RNA Cap Structure Analog (10 μmol) (New England BioLabs, Ipswich, MA), 2 µg linearized DNA, 2 µl RNaseOUT™ RNase inhibitor (Invitrogen, Carlsbad, CA). After 2 h incubation at 37 °C, DNA templates were eliminated by treatment with 2 µl TURBO™ DNase (Invitrogen, Carlsbad, CA) for 30 min at 37 °C. *In vitro* transcribed RNAs were purified by NucAway™ Spin Columns (Invitrogen, Carlsbad, CA), and RNA concentration was quantified by spectrophotometer-SimpliNano (GE Healthcare Life Sciences, Pittsburgh, PA).

### Genomic and subgenomic RNA quantification

For intracellular viral RNA quantification, BHK-21 cells were seeded on 24-well plates at a density of 1 × 10^5^ cells/well 24 h before transfection. *In vitro* transcribed viral genomic RNA (1 µg) was transfected into BHK-21 cells using Lipofectamine™ MessengerMAX™ Transfection Reagent (Invitrogen, Carlsbad, CA). Cell culture supernatants were harvested at 18 and 36 h post transfection (hpt). Experiments were performed in triplicate at each time point. At 18 hpt, cells were lysed using RNA Lysis Buffer of SV Total RNA Isolation System (Promega, Madison, WI) for RNA quantification.

To precisely quantify the ratios of gRNA and sgRNA (plus strand or minus strand), RT-qPCR was conducted with a panel of specific primers/probe set, which were designed to cover the leader-B-TRS-junction sites as we described previously (30). For a nucleotide sequence in the TRS of sgRNA2, 6, and 7 that shown heterogeneity, a degenerate nucleotide was used in the probes (30). Primer/probe sets were synthesized by Biosearch Technologies (Petaluma, CA).

To conduct the qRT-PCR test, initially, total RNA was extracted from BHK-21 cells transfected with viral genomic RNAs by using the SV Total RNA Isolation System (Promega, Madison, WI) based on the user’s manual. Extracted RNAs were also quantified by SimpliNano (GE Healthcare Life Sciences, Pittsburgh, PA). Total RNA (500 ng) was reverse transcribed by Maxima H Minus Reverse Transcriptase (Thermo Scientific, Carlsbad, CA). Plus-strand genomic RNA, plus-strand subgenomic RNA, and minus-strand genomic/subgenomic RNA were reverse transcribed in three different reactions by using three different sets of primers (30). Real-time PCR was performed in 20 µl reactions using 10 µl TaqMan™ Fast Advanced Master Mix (Applied Biosystems, Carlsbad, CA), 0.25 µl forward primer (40 µM), 0.25 µl reverse primer (40 µM), 0.4 µl Taqman probe (10 µM), 2 µl cDNA, 7.1 µl H_2_O. PCR reactions were performed on a CFX96 Touch™ Real-Time PCR Detection System (Bio-Rad, Hercules, CA). Conditions were set up based on the TaqMan™ Fast Advanced Master Mix user’s instructions: 50 °C UNG incubation 2 min, 95 °C polymerase activation 2 min, then 40 cycles of 95 °C denaturation 3 s, 60 °C annealing / extention 30 s. The relative accumulation ratio of sgRNAs was quantified using the threshold cycle (_ΔΔ_CT) method (28, 29). To precisely quantify RNA accumulation and to rule out possible background interference from initial transfected RNA, a transcription-defective mutant was constructed, in which the nucleotide sequence from 495 nt to 515 nt was replaced with 7 consecutive stop codons to terminate the translation of nonstructural polyproteins. The (+) gRNA detected from BHK-21 cells transfected with (+) gRNA of the defective mutant was taken as the background level. In the quantification involved in (+) gRNA, the input genomic RNA level derived from transfection was considered as background and deducted at first.

### Subgenomic RNA accumulation ratio deviation

For each (+) sgRNA to (+) gRNA ratio *r_virus_*, *_sgRNA_* (*virus* = WT or one of the mutants, *sgRNA* = sgRNA 2, 6 or 7), the mean value of *r_virus_*, *_sgRNA_* was first obtained by averaging values of 3 replicates of each virus and standard deviation *s_virus_*, *_sgRNA_* was calculated based on all values of *r_virus_*, *_sgRNA_*. Next, for each mutant and each sgRNA, the deviation of *r_mutant_*, *_sgRNA_* from *r_WT_*_,*sgRNA*_ was calculated as a *z*-score *z_mutant_*_, *sgRNA*_ = (*r_WT_*_, *sgRNA*_ – *r_mutant_*_, *sgRNA*_) / *s_virus_*_, *sgRNA*_. For a given mutant, the *z*-score values for sgRNAs 2, 6 and 7 were then averaged to obtain the mean deviation of sgRNA accumulation ratio z_J*_mutant_* = (*z_mutant_*, *_sgRNA2_* + *z_mutant_*, *_sgRNA6_* + *z_mutant_*_, *sgRNA7*_)/3.

### Virus titration

Supernatants from transfected BHK-21 cells were harvested at 18 hpt and 36 hpt, then titrated on MARC-145 cells using the TCID_50_ method as previously described (28). For more precise quantification, the immunofluorescence assay (IFA) was used to facilitate TCID_50_ quantification. At 96 h post infection, MARC-145 cells in 96-well plates were fixed with ice-cold methanol for 30 mins at minus −20 °C, then allowed to air dry. The anti-N protein monoclonal antibody-SDOW17 was used as the primary antibody. Plates were incubated at 37 °C for 1 h with SDOW17 diluted in phosphate-buffered saline (PBS), then washed three times with PBS. Alexa Fluor 488 AffiniPure donkey anti-mouse IgG (H+L) (Jackson Immuno Research, West Grove, PA) was used as secondary antibody. After 1 h incubation at 37 °C with secondary antibody, plates were washed three times, then the plate was read using a fluorescence microscope EVOS FL Cell Imaging System (Thermo Fisher Scientific, Carlsbad, CA).

## RESULTS

### Identification of an intra-family conserved high-order RNA structure in the M ORF of arteriviral genomes

To identify novel functional elements in PRRSV genomes, we applied synplot2 (17) to alignments of ∼200 PRRSV-1 and ∼200 PRRSV-2 sequences (see Methods). Synplot2 analyses the relative levels of synonymous substitutions at different sites in protein-coding sequences. Regions where there is a statistically significant deficit of synonymous substitutions relative to the genome average are indicative of overlapping functional elements (17). As shown in Fig. 1A, for both PRRSV-1 and PRRSV-2, strong and statistically significant peaks in synonymous site conservation were found coinciding with the TF ORF that overlaps ORF1a, the E ORF that overlaps GP2, the 5a ORF that overlaps GP5, the dual coding regions where GP3 and GP4 overlap and where GP2 and GP3 overlap, and the ribosomal frameshift signal (a U_UUA_AAC shift site and a 3′-adjacent RNA pseudoknot structure) at the junction of ORFs 1a and 1b. Additional synonymous site conservation peaks were observed near the 5′ end of ORF1a, throughout the N ORF, just upstream of GP5 and, in PRRSV-2, just upstream of GP3 and at the junction of GP5 and M. In addition, a particularly striking extended region of synonymous site conservation was observed to cover the 3′ half of the M ORF, in both PRRSV-1 and PRRSV-2. To our knowledge, this feature cannot be explained by currently known functional elements in PRRSV genomes.

Fig. 1B shows an expanded view of the M and N ORF region of the PRRSV-1 and PRRSV-2 synonymous site conservation plots. RNA structure predictions for this region for representative sequences of PRRSV-1 and PRRSV-2 are shown in Fig. 2. Despite substantial sequence divergence between PRRSV-1 and PRRSV-2, many elements of the structure prediction are conserved between the two – including two stem-loops (SL1 and SL2 on Fig. 2) and base pairing between two more distal regions (extended stem-loop, extSL on Fig. 2). We decided to focus on these elements as they also correspond to some of the regions of highest synonymous site conservation (coloured horizontal bars in Fig. 1B: yellow – SL1, orange – SL2, green – two halves of the extSL duplex). Although PRRSV-1 and PRRSV-2 differ in primary sequence (∼70% nucleotide identity), the SL1, SL2 and extSL structures were predicted in both the PRRSV-1 and PRRSV-2 sequences analysed, with three pairs of fully compensatory substitutions (e.g. C:G to U:A) in each of SL1 and SL2, besides one partially compensatory substitution (i.e. single-nucleotide changes between a G:U base pair and a G:C or A:U base pair) in each of SL1, SL2 and extSL.

**Figure 2.**
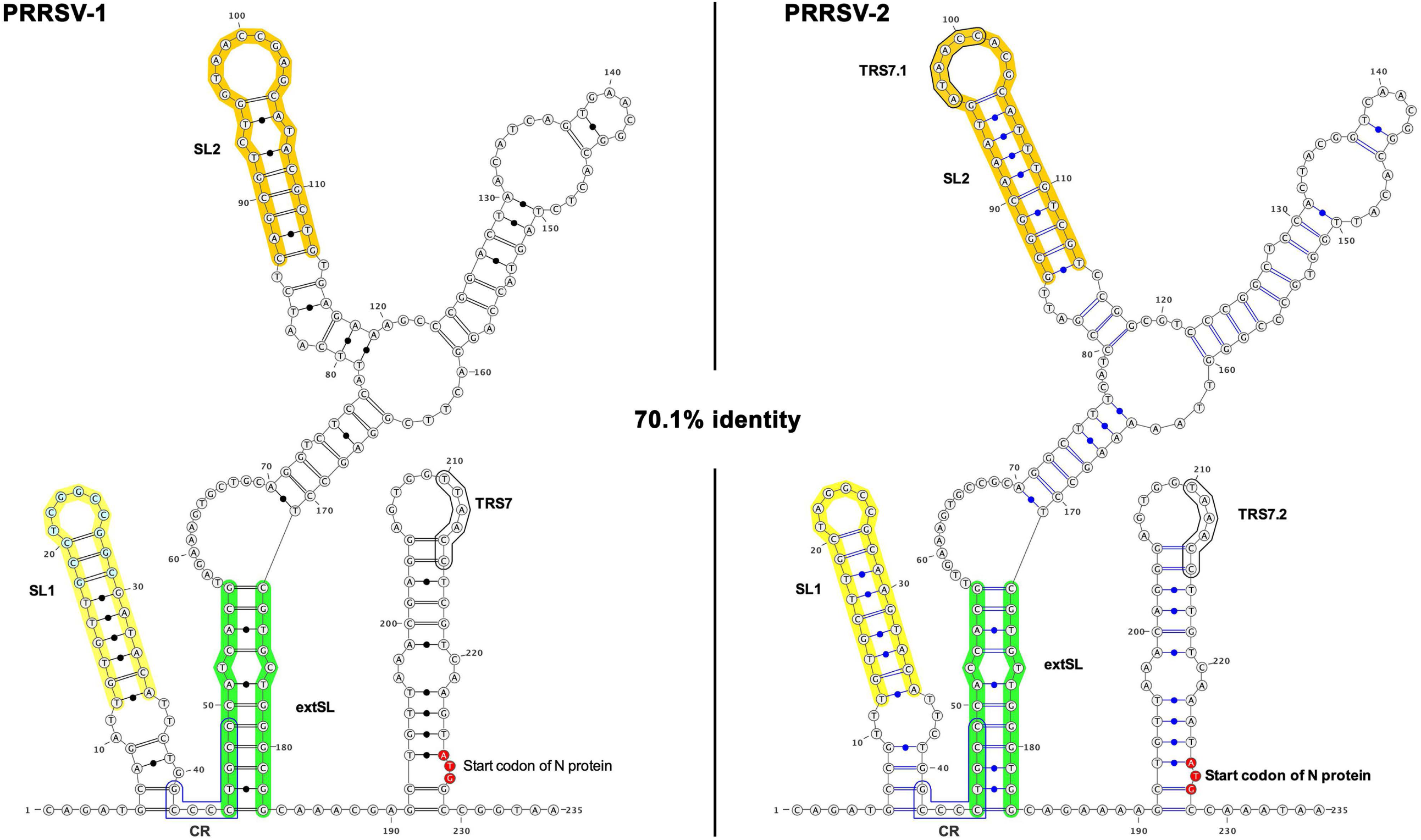
RNA structure prediction of the synonymously conserved region within PRRSV M ORF. RNA structure predictions for the 3′ half of the M ORF and beginning of the N ORF regions for the PRRSV-1 (14323-14557nt, DQ489311.1) and PRRSV-2 (14666-14900 nt, KC469618.1) sequences. Minimum free energy structures were predicted with RNAstructure (21). The SL1, SL2 and extSL duplex regions are annotated, as well as an additional conserved region (CR), the transcription regulatory sequence for the N protein sgRNA 7 (TRS7), an additional sgRNA 7 TRS in PRRSV-2 (TRS7.1), and the N ORF initiation codon.

To test further for conservation of structure, we extracted the sequences corresponding to the SL1, SL2, and extSL duplexes from 461 PRRSV-1 and 1518 PRRSV-2 genome sequences (see Methods). All unique sequences, and the number of times that each unique sequence was found, for each of SL1, SL2 and extSL are shown in Fig. 3. SL1 is supported by seven different pairs of fully compensatory substitutions, while single mispairings were observed in only 21 of the 1979 sequences. The primary sequence of extSL is in general highly conserved between PRRSV-1 and PRRSV-2; nonetheless, the functional importance of the predicted duplex was supported by four distinct pairs of fully compensatory substitutions when all 1979 sequences were considered. A single-nucleotide paired bulge (either U.C or C.U) is present in extSL in every sequence (Fig. 3, green arrows) but, aside from this ubiquitous bulge, spurious mispairings were again present in only a handful of sequences (14 of 1979). SL2 is relatively less conserved. A mispairing at the 7th position from the base of SL2 was ubiquitous in PRRSV-1 sequences (Fig. 3, yellow arrows), whereas in PRRSV-2 the duplex was generally uninterrupted. In addition, 22 of 461 PRRSV-1 and 46 of 1518 PRRSV-2 sequences have other mispairings in SL2, sometimes with multiple mispairings in the same sequence. Nonetheless, SL2 is uninterrupted in 97% of PRRSV-2 sequences and – aside from the aforementioned single mispairing at the 7th position – otherwise uninterrupted in 95% of PRRSV-1 sequences; it is also supported by six pairs of fully compensatory substitutions across the 1979 sequences.

**Figure 3.**
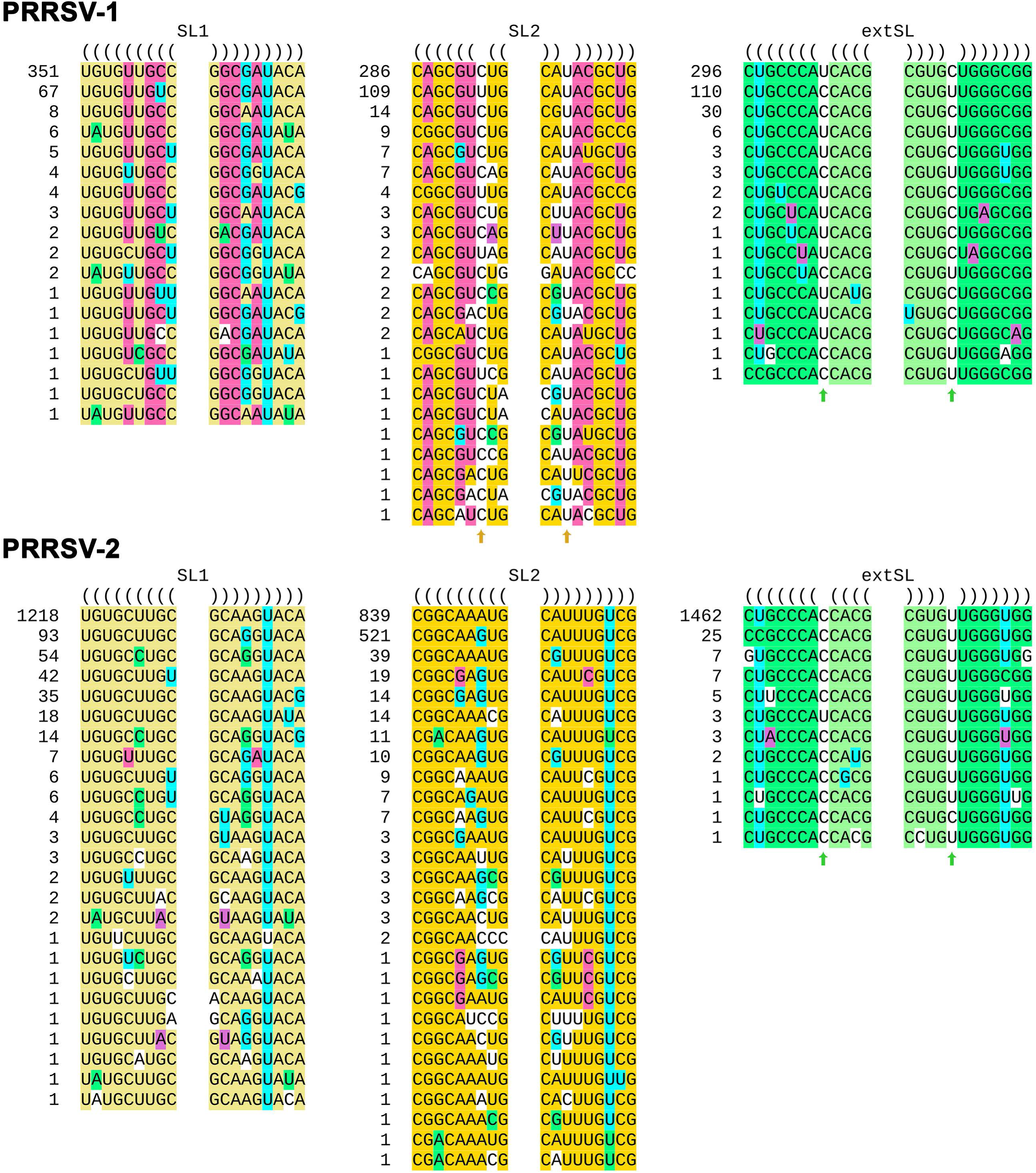
Analysis of the SL1, SL2 and extSL duplexes in PRRSV-1 and PRRSV-2 genome sequences. Each unique sequence was shown with the number at left indicating how many times that unique sequence occurs among the 461 or 1518 sequences. One PRRSV-1 extSL sequence was omitted as it contained an ambiguous nucleotide code (viz. "R"). Predicted base pairings were highlighted in yellow (SL1), orange (SL2) or green (extSL), except that G:U base pairs are indicated by highlighting either the G or the U in cyan. Compensatory substitutions were highlighted in magenta, green, or purple. Single-nucleotide paired bulges (either U:C or C:U) present in SL2 and extSL were indicated with arrows. Unpaired nucleotides are not highlighted.

PRRSV-1 and PRRSV-2 do not on their own form a monophyletic clade, but instead they are separated by various arteriviruses that have been isolated from rodents in phylogenetic trees (31, 32). PRRSVs and several rodent arteriviruses were classified into the genus *Betaarterivirus*, subfamily *Variarterivirinae*. Thus, we inspected the sequences of other members of this clade (Genera *Betaarterivirus* and *Nuarterivirus* of subfamily *Variarterivirina* demarcated by the red circle in Fig. S1;. A single sequence NC_075972 representing *Nuarterivirus*.) to see whether they might also have predicted SL1, SL2, and extSL duplexes. Once again, we found that SL1 and extSL were well conserved whereas SL2 was more variable between sequences and more often contained multiple mispairings (Fig. 4A). We analysed the individual sequences with RNAfold (20), which predicts minimum free energy RNA structures of single sequences. When applied to the sequence region shown in Fig. 4A, RNAfold predicted SL1 and extSL for all seven sequences, whereas SL2 was predicted in all sequences except NC_032987 where it was disrupted by competing alternative structure. We also inspected these sequences with RNAz which analyses structural conservation and thermodynamic stability to identify statistically significant conservation of RNA structure in sequence alignments (33). With the default 120 nt sliding window and 40 nt step size, a 120 nt window enclosing SL1 (but not SL2 or extSL) was the highest scoring window in the whole genome (Fig. 4B).

**Figure 4.**
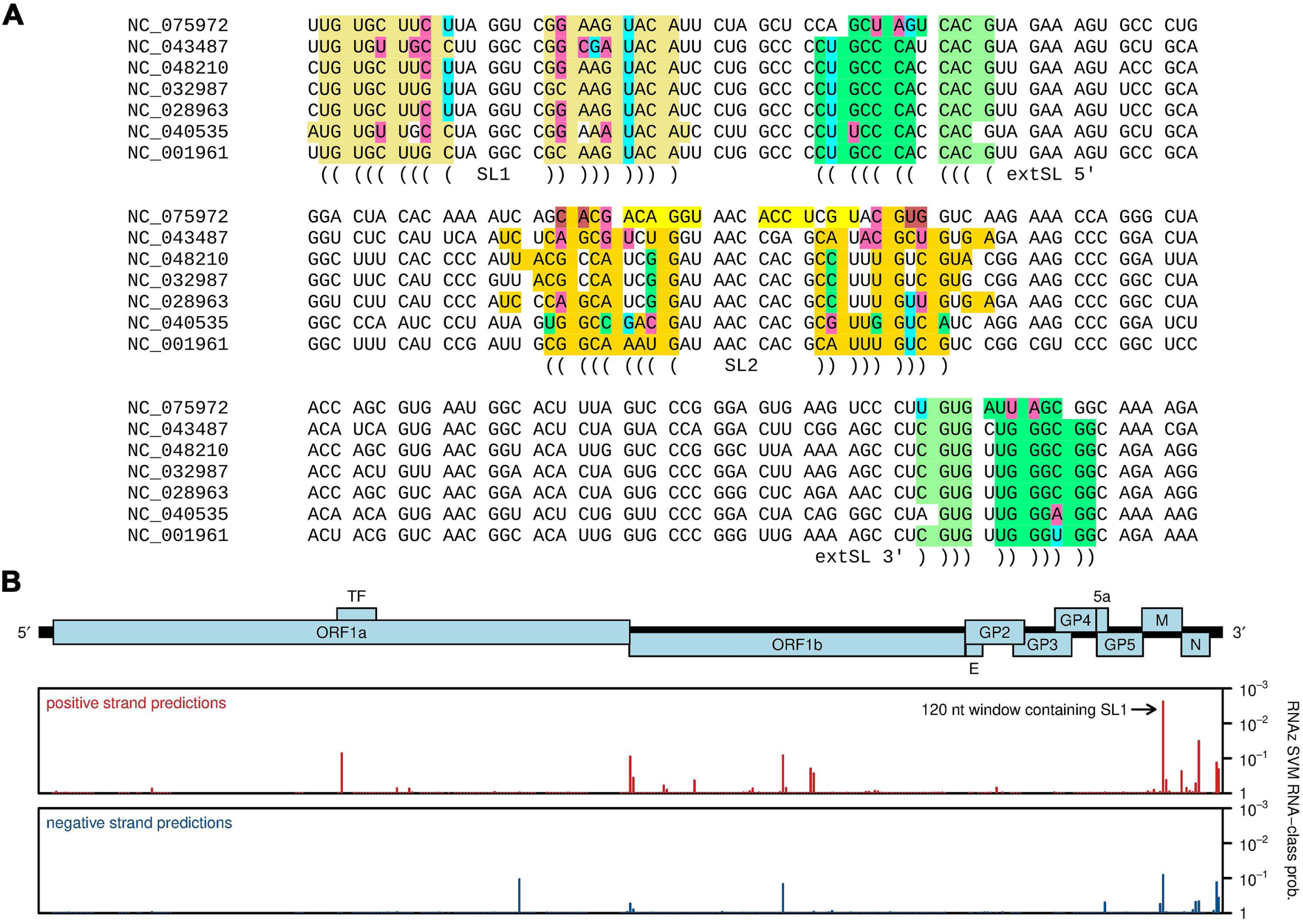
Conservation of SL1, SL2 and extSL across genera *Betaarterivirus* and *Nuarterivirus*. **(A)** The SL1-SL2-extSL region of NCBI RefSeqs in the *Betaarterivirus*/*Nuarterivirus* clade with predicted structures annotated using a color scheme similar to that used in Figure 3. M ORF codons are separated by spaces. **(B)** RNAz analysis of an alignment of the six RefSeqs NC_001961 (PRRSV-2), NC_028963 (Rat arterivirus 1), NC_040535 (Rodent arterivirus), NC_043487 (PRRSV-1), NC_048210 (Rodent arterivirus) and NC_075972 (Rodent arterivirus). The *y*-axis plots 1 − *p*, where *p* is the RNAz SVM RNA-class probability statistic. Bars correspond to the midpoints of 120 nt windows, mapped from alignment coordinates (RNAz output) back to NC_001961 (PRRSV-2) coordinates.

Similar RNA structures at equivalent positions were also predicted in other arteriviruses. For a clade containing lactate dehydrogenase-elevating virus (LDV) (genus *Gammaarterivirus* and relatives; green text in Fig. S1), SL1, SL2, and extSL structures were predicted manually in all inspected viruses except the basally branching Lopma virus (MW595222) (Fig. S2). For a clade containing simian hemorrhagic fever virus (SHFV) (subfamily *Simarterivirinae*; blue text in Fig. S1), SL1 and extSL but not SL2 were predicted manually in all inspected viruses (Fig. S3). As well as phylogenetic conservation, these manual predictions were also supported by a number of compensatory substitutions (Fig. S2 and Fig. S3). With the exception of Lopma virus, the RNAfold minimum free energy fold of the displayed sequence regions also supported SL1 in 5 of 7 sequences, SL2 in 7 of 7 sequences and extSL in 6 of 7 sequences for the "LDV" clade, and both SL1 and extSL in all 16 sequences of the "SHFV" clade (though with minor differences in the 5′ and 3′ extent of some duplexes due to terminal G:U base pairs and/or alternative, equally optimal base pairings).

### Disruption of PRRSV M ORF RNA structure impairs infectious virus production

To investigate the potential role of the M ORF RNA structure in viral replication, synonymous mutations were designed to disrupt the structure in the genome of a PRRSV-1 isolate SD01-08 (Fig. 5). For mutants of stem-loop SL1, 5SL1, and 3SL1 constructs contain complementary mutations disrupting the 5′ side and 3′ side of SL1, respectively, while Loop1 mutations alter the loop sequence and 53SL1 contains both mutations from 5SL1 and 3SL1 to restore the stem structure (Fig. 5A). A similar design was applied to SL2, generating four mutants – 5SL2, 3SL2, Loop2, and 53SL2 (Fig. 5B). For extSL, mutants were initially designed based on mutating highly conserved codon positions identified from the synplot2 analysis. The conserved region (CR) mutant contains three synonymous mutations that disrupt the 5′ side of the extSL duplex and also the preceding codon, whereas the extSL3 mutant contains four synonymous mutations that disrupt the 3′ side of the extSL duplex (Fig. 5C). Mutations in CR may only partially weaken the 5′ side of extSL.

**Figure 5.**
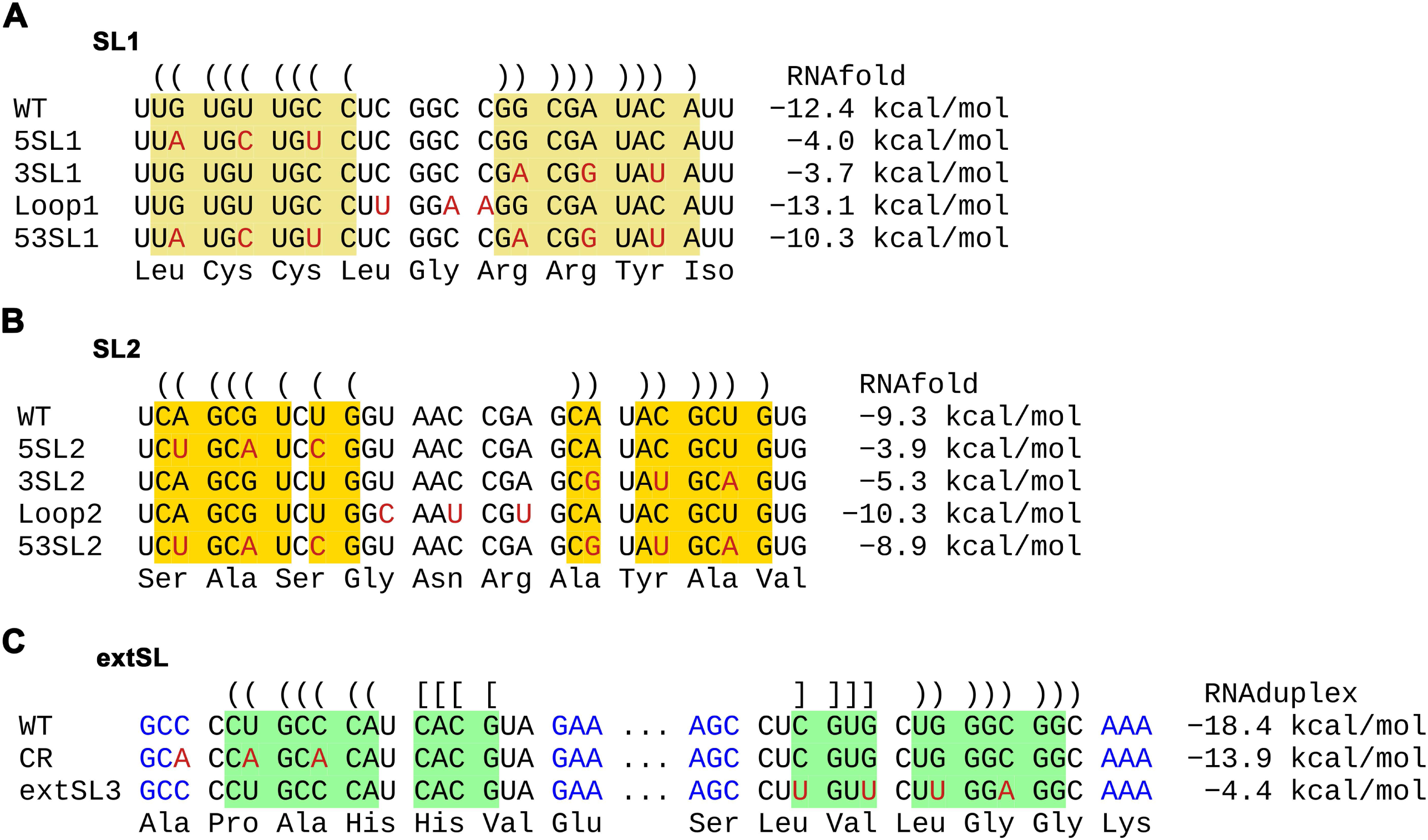
Design of mutagenesis for the high-order RNA structure in the M ORF of PRRSV-1. Synonymous mutations introduced in stem-loop1 **(A)**, stem-loop2 **(B)**, and extSL **(C)**. Nucleotides forming stem1, stem2 and extSL are highlighted in yellow, orange and green, respectively. Parenthesis or brackets are used to indicate predicted base pairings in WT. The free energy of each structure after mutation is shown at the right side of each sequence.

Initially, we determined the impact of these mutations on infectious virus production. Genomic RNAs of wild-type (WT) and mutants were *in vitro* transcribed and then transfected into BHK-21 cells. At 18 and 36 hours post-transfection (hpt), viral titer in the supernatant of transfected cells was quantified (Fig. 6). Infectious viruses of 5SL1 and 3SL1 could not be detected at 18 hpt (Fig. 6A). At 36 hpt, viral titers for 5SL1 and 3SL1 were reduced 2.5- and 2.1-log, respectively, compared with WT (Fig. 6B). The mutations introduced in the Loop1 mutant had less impact on viral production with the viral titer decreased by only 0.7- and 0.8-log compared to WT at 18 hpt and 36 hpt, respectively. SL2 seems to be less critical than SL1 for viral production. At 18 hpt and 36 hpt, viral titers for the 5SL2 mutant were reduced 1.2- and 1.7-log, respectively, while viral titers for the 3SL2 mutant were decreased 0.6- and 1.1-log. There was no significant difference in viral titer between WT and the Loop2 mutant (Fig. 6A and 6B).

**Figure 6.**
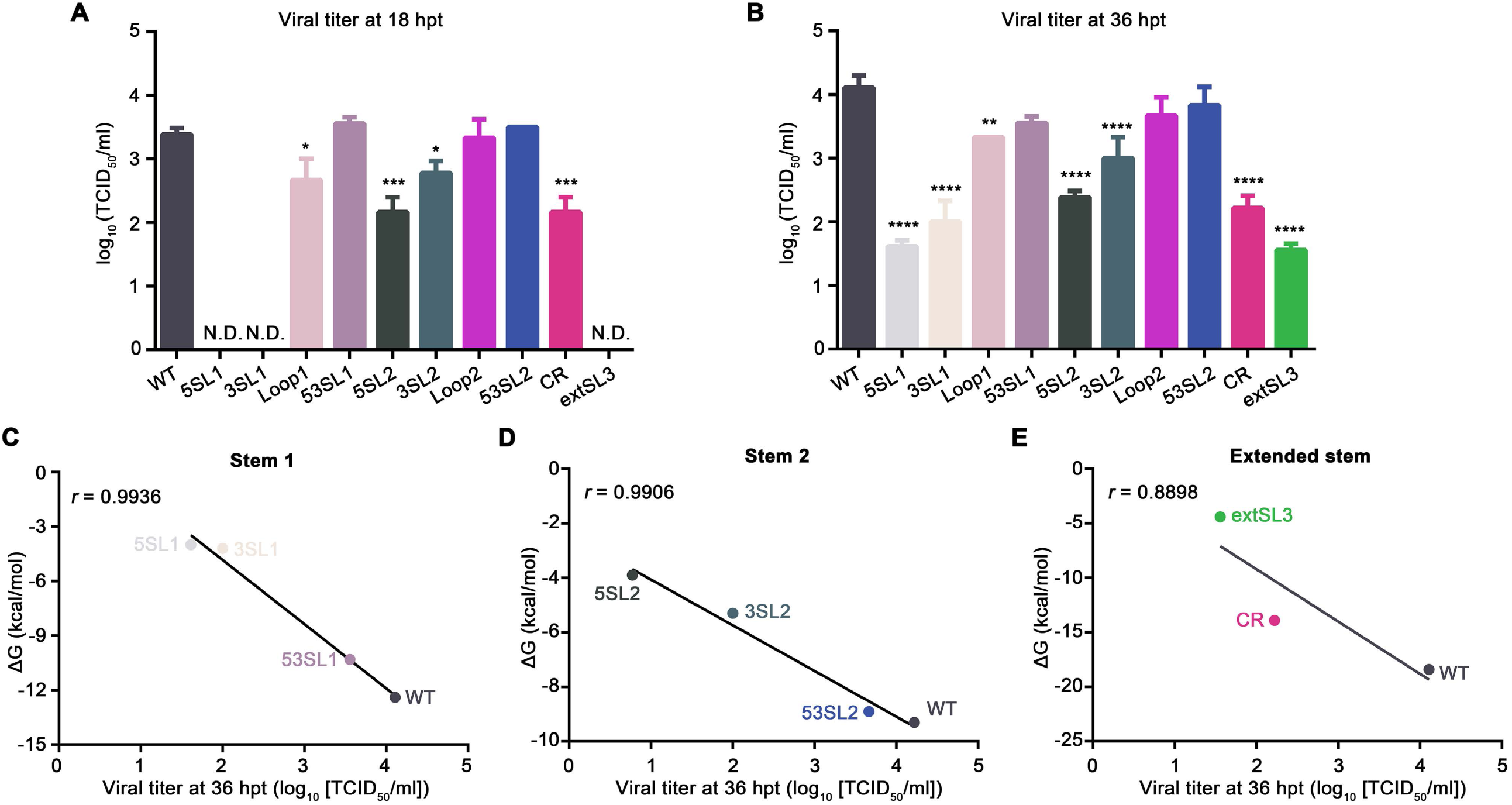
Mutations disrupting the conserved RNA structure within the PRRSV-1 M ORF decrease the level of recombinant virus production. (A-B) BHK-21 cells were transfected with *in vitro*-transcribed gRNA and recombinant viruses were harvested at 18 hpt **(A)** and 36 hpt **(B).** Virus titers were quantified on MARC-145 cells using the TCID_50_ method. Each sample was tested in triplicate. The titers of mutant viruses were compared with that of WT virus by one-way analysis of variance (ANOVA) followed by Tukey’s post hoc test. *: *P* < 0.05, **: *P* < 0.01, ***: *P* < 0.001, ****: *P* < 0.0001. **(C-E)** Pearson correlation between the estimated free energy of structures and viral titers for stem 1 mutants **(C)**, stem 2 mutants **(D)**, and extSL mutants **(E)**. The titers of the mutant viruses at 36 hpt were plotted against the corresponding free energy of the mutated stem structures. A negative linear regression is depicted as a black line. The Pearson correlation coefficients (*r*) are shown in the corresponding plots.

As expected, mutations designed to restore the SL1 and SL2 duplexes, 53SL1 and 53SL2, were able to recover the viral production, with no significant difference observed in viral titer between WT and these two mutants (Fig. 6A and 6B). Mutations in CR and extSL3 both showed significant impact on viral production. The titer of CR was decreased by 1.2- and 1.9-log at 18 and 36 hpt, respectively (Fig. 6A and 6B). Mutant extSL3 – whose mutations may theoretically more seriously disrupt extSL than those of mutant CR – was consistently more attenuated. At 18 hpt, no infectious viruses were detected (Fig. 6A); whereas at 36 hpt, the titer of the extSL3 mutant was decreased by 2.6-log in comparison to that of WT (Fig. 6B).

The thermodynamic stability of stem-loop structures in mutant viruses was also associated with the viral yield. Statistical analysis revealed that the free energy of all viral constructs in the three stems within the high-order RNA structure correlated well with the level of infectious virus production [Pearson correlation coefficient (r) > 0.99 in Fig. 6C-D and less extent (r > 0.89) in Fig. 6E]. Mutants 5SL1 and 3SL1 with high free energy of SL1 showed the lowest viral titers (Fig. 6C). As expected, 53SL1 designed to maintain the thermo-stability of SL1 could restore the level of viral production (Fig. 6C). The free energy changes of SL2 were also correlated well with the changes of viral titer (Fig. 6D). Mutant 5SL2 with the highest free energy had the lowest virus yield, while mutant 3SL2 with less free energy loss than its complimentary 5SL2 showed less reduction of viral titer (Fig. 6D). Similar to 53SL1, 53SL2 that was designed to restore the SL2 duplex, had a free energy level and viral titer close to that of WT virus. Viral yield of extSL mutants is also statistically correlated with the stability of its RNA structure (Fig. 6E). In summary, synonymous mutations destabilizing the high-order RNA structure decrease infectious virus production, in which SL1 and extSL play a more important role.

### The PRRSV M ORF RNA structure regulates the relative accumulation levels of viral subgenomic RNAs

For coronavirus subgenomic RNA (sgRNA) transcription, two long-distance RNA-RNA interactions have been described with critical roles in facilitating the discontinuous transcription of the most abundant, N-protein-coding sgRNAs of TGEV (5, 7). In the predicted PRRSV M ORF high-order RNA structure model, the TRS7 for sgRNA 7 (encoding the N protein) is located immediately downstream of extSL, which led us to hypothesize that arteriviruses might also use a similar strategy to enhance the discontinuous transcription of sgRNAs. We therefore quantified the relative accumulation ratio of sgRNAs to genomic RNA (gRNA) using RT-qPCR (Fig. 7A). Specifically, the relative abundances of sgRNA 6 (encoding the M protein) and sgRNA 7 to gRNA were analyzed. We hypothesized that the transcription of sgRNA 2 (encoding the GP2 and E proteins) – the sgRNA with the body (B)-TRS most distal from the M ORF RNA structure – should be least affected by any modulatory effects of the structure. Therefore, the ratio of sgRNA 2 to gRNA was also quantified as a control.

**Figure 7.**
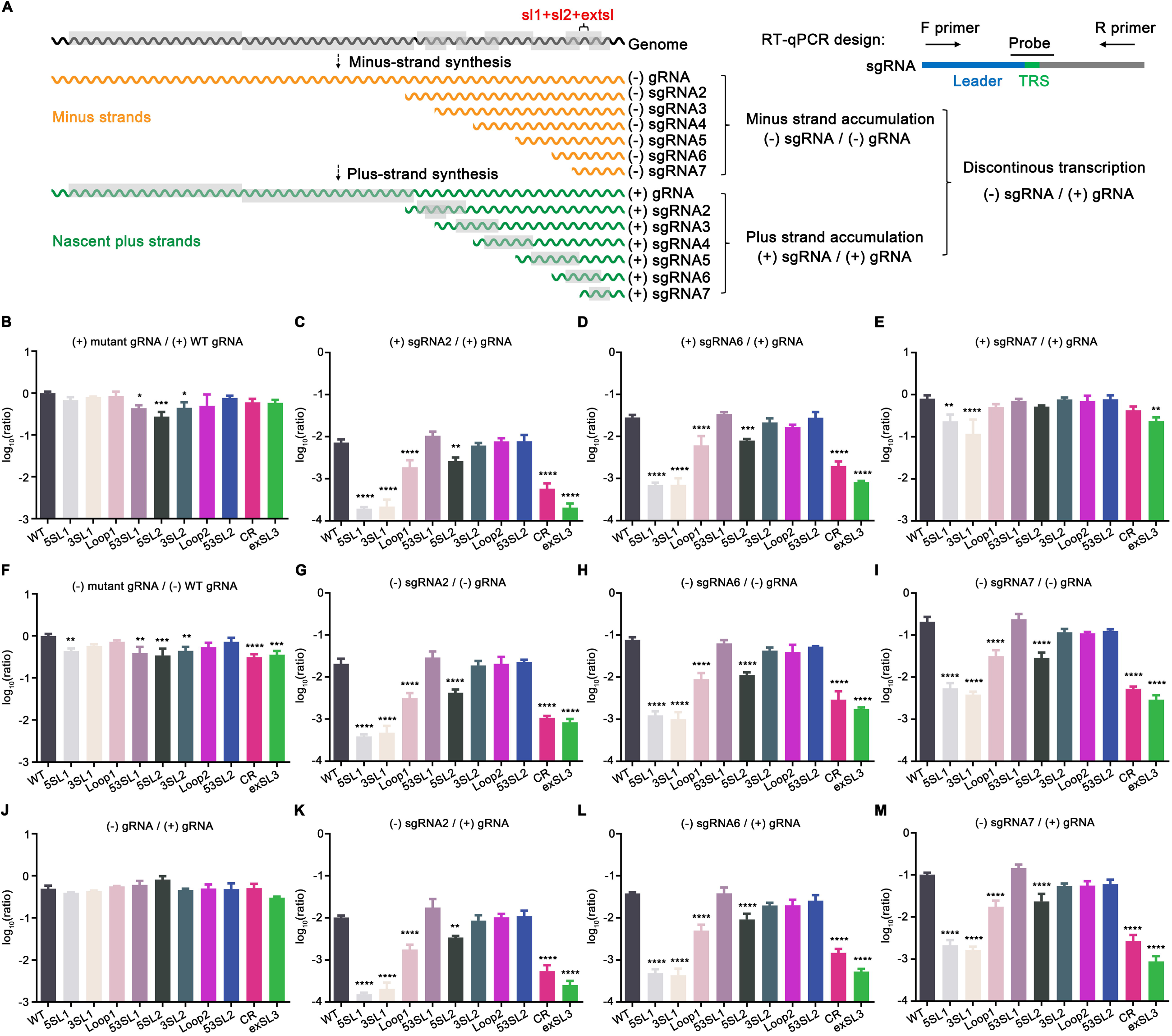
Effect of M ORF stem-loop mutations on the accumulation levels of (+) and (−) genomic and subgenomic RNA 2, 6 and 7. **(A)** Diagram of arteriviral RNA transcription and replication. (B-M) *In vitro* transcribed gRNA of WT or mutant viruses were transfected into BHK-21 cells. Viral RNAs were quantified at 18 hpt. The level of viral RNAs was normalized to host housekeeping gene (TBP) first before comparing with other viral RNAs. Each sample was tested in triplicate. (**B**) Ratio of mutant (+) gRNA to WT (+) gRNA. (**C-E**) Ratio of (+) sgRNA 2/6/7 to (+) gRNA for each virus. (**F**) Ratio of mutant (-) gRNA to WT (-) gRNA. (**G-I**) Ratio of (-) sgRNA 2/6/7 to (-) gRNA for each virus. (**J**) Ratio of mutant (-) gRNA to WT (+) gRNA. (**K-M**) Ratio of (-) sgRNA 2/6/7 to (+) gRNA for each virus. Statistical differences were evaluated by one-way analysis of variance (ANOVA) followed by Tukey’s post hoc test. *: *P* < 0.05, **: *P* < 0.01, ***: *P* < 0.001, ****: *P* < 0.0001.

BHK-21 cells were transfected with gRNA of WT or mutant viruses, harvested at 18 hpt, and the relative accumulation of gRNA and sgRNAs were quantified. Initially, we analyzed the relative accumulation ratio of (+) sgRNAs to the corresponding (+) gRNA, as well as the relative ratio of mutant (+) gRNAs to WT (+) gRNA (Fig. 7B-E).

Surprisingly, results showed that the relative ratio of the three subgenomic RNAs – sgRNA 2, sgRNA 6, and sgRNA 7 – were all affected by mutations disrupting the M ORF RNA structure. In comparison with that of WT virus, the relative abundance of (+) sgRNA 2 to (+) gRNA was severely reduced by 37.8, 32.1, 3.7, 12.3, and 34.9 folds in mutants 5SL1, 3SL1, Loop1, CR, and extSL3, respectively. For mutant 5SL2, the relative accumulation of (+) sgRNA 2 to (+) gRNA was also decreased by 2.8-fold compared to WT. As expected, the (+) sgRNA 2 to (+) gRNA was restored for the 53SL1 and 53SL2 mutants. The relative accumulation of (+) sgRNA 6 to (+) gRNA was also affected in a similar pattern to that seen for (+) sgRNA 2. The loss of relative accumulation of (+) sgRNA 7 to (+) gRNA was much less severe and only significant for mutants 5SL1, 3SL1, CR, and extSL3, in which it was reduced by 3.3, 5.8, 1.9, and 3.4 folds, respectively. In contrast, we observed only slight fluctuation of (+) gRNA accumulation, which seems to be not correlated with the loss of sgRNAs accumulation. The four mutants – 5SL1, 3SL1, CR, and extSL3 – that had greater impact on the ratios of sgRNA accumulation, did not exhibit a similar impact on gRNA accumulation, suggesting that the conserved high-order RNA structure may mainly function in sgRNA accumulation instead of gRNA accumulation, in which the SL1 and extSL may play a more important role.

Based on the current model of arterivirus RNA synthesis (34), the abundances of continuously transcribed (−) gRNA and discontinuously transcribed (−) sgRNAs are believed to determine the accumulation levels of (+) gRNA and (+) sgRNAs. Therefore, we also calculated the relative ratios of (−) sgRNAs to their corresponding (−) gRNA, and the relative ratios of mutant (−) gRNAs to WT (−) gRNA (Fig. 7F-I). We only observed a modest decrease of the level of (−) gRNA in mutant gRNA-transfected cells. In contrast, (−) sgRNA 2, 6, and 7 to (−) gRNA ratios of the mutants showed a largely consistent pattern with that of the plus strand RNAs as shown in Fig. 7C-E.

Since plus-strand gRNAs function as direct templates for discontinuous and continuous transcription, the ratios of mutant (−) gRNA and (−) sgRNAs to their corresponding (+) gRNAs were also calculated (Fig. 7J-M). Consistently, no significant changes were observed in the (−) gRNA / (+) gRNA ratio among these mutants, but the changes in relative accumulation ratio of (−) sgRNA 2, 6, and 7 to (+) gRNA showed a similar pattern as that presented in Fig. 7C-E and 7G-I.

### Accumulation levels of subgenomic RNAs correlate with the production of viral progeny

Since arterivirus structural proteins are translated from (+) sgRNAs, the yield of viral particles will theoretically correlate with the production of (+) sgRNAs (34). To further investigate the correlation between the change of subgenomic RNA levels and recombinant virus yields, the data obtained on the relative ratios of (+) sgRNA2/6/7 to (+) gRNA at 36 hpt were plotted against the titers of the WT and mutant viruses (Fig. 8A-E). The modest fluctuation of (+) gRNA accumulation levels in WT and mutant viruses was not correlated with infectious virus production (*r* = 0.36), suggesting the decrease of mutant virus production is not determined by the change of (+) gRNA accumulation (Fig. 8A). As expected, the decreased ratios of (+) sgRNA 2/6/7 to (+) gRNA were well correlated with the decreased titers of viruses (*r* > 0.85) (Fig. 8B-D). The five mutants (5SL1, 3SL1, 5SL2, CR and extSL3) with significantly reduced ratios of subgenomic RNA accumulation correlated well with significantly decreased levels of viral production. We also calculated the mean deviations of the (+) sgRNA to (+) gRNA ratios (comparing mutant with WT ratios and averaging across the 3 sgRNAs; see Methods for details).

**Figure 8.**
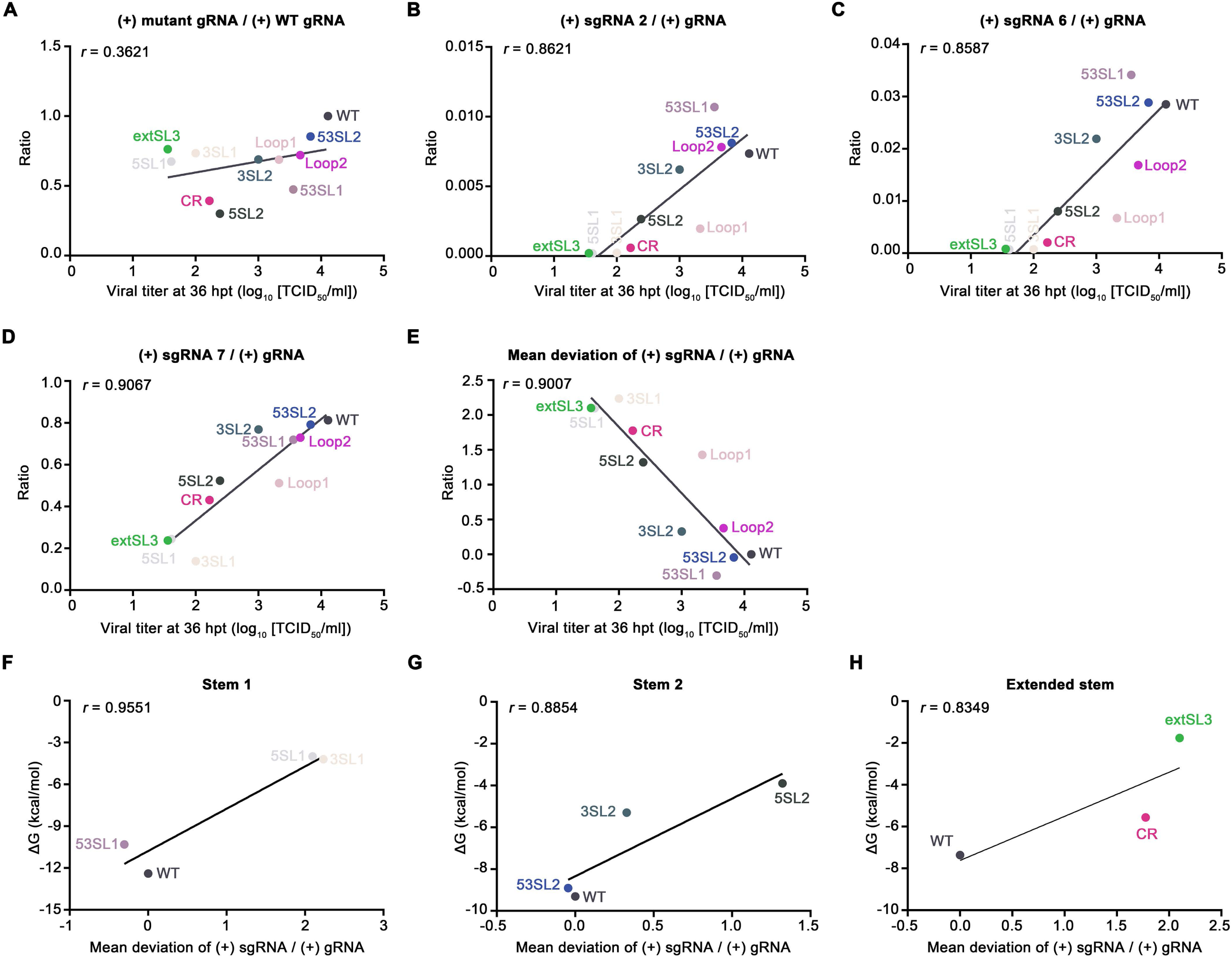
Correlations between relative accumulation of sgRNAs and infectious virus production or thermodynamic stability of M ORF structures. **(A)** Virus titer at 36 hpt compared with mutant (+) gRNA to WT (+) gRNA ratio. **(B-D)** Virus titer at 36 hpt compared with (+) sgRNA 2 to (+) gRNA ratio **(B)**, (+) sgRNA 6 to (+) gRNA ratio **(C)**, and (+) sgRNA 7 to (+) gRNA ratio **(D)**. **(E)** Virus titer at 36 hpt compared with mean deviation of sgRNA ratio from WT. **(F-H)** Mean deviation of sgRNA ratio from WT compared with free energy of stem 1 mutants **(F)**, stem 2 mutants **(G)**, or extSL mutants Data fitting linear regressions are depicted as black lines. The corresponding Pearson correlation coefficient value (*r*) is shown for each plot.

Mutants 53SL1, 53SL2, and Loop2 with mean deviations from WT close to zero (WT virus phenotype) produced a similar level of infectious particles as WT virus (Fig. 8E). In contrast, for those mutants 5SL1, 3SL1, 5SL2, CR, and extSL3 that had large mean deviations from WT in sgRNA accumulation, significantly lower viral titers were observed (Fig. 8E). A high Pearson correlation coefficient (*r* = 0.9007) was obtained, which demonstrates that the loss of accumulation of sgRNAs in the mutants is closely correlated with decreased infectious particle production.

Consistent with the correlation between thermodynamic stability of stem structures and the virus titer as shown in Fig. 6C-E, the free energy changes of stem structures were also correlated with the mean deviation from WT of sgRNA accumulation (*r* > 0.83; Fig. 8F-H). The mutants (5SL1, 3SL1, 5SL2, CR, and extSL3) with higher free energy showed greater loss of sgRNA accumulation. In contrast, mutants 53SL1 and 53SL2 that were designed to repair the stability of stems could successfully restore sgRNA accumulation (Fig. 8F-H). Collectively, these data suggest that the thermodynamic stability of the high-order RNA structure in M ORF is correlated with the relative accumulation of sgRNAs.

## DISCUSSION

In nidoviruses, the discontinuous transcription of sgRNAs is precisely regulated by both *cis*-acting and *trans*-acting factors (3–7). To our knowledge, the current study is the first to identify a *cis*-acting element within protein-coding regions that is able to modulate arterivirus sgRNA transcription. We initially thought that the conserved RNA structure might functionally resemble the *cis*-acting element in TGEV that enhances the discontinuous transcription of the most abundant sgRNA – that expressing the N protein (5, 7). In this model, the structure would attenuate RTC processivity in the vicinity of the N ORF B-TRS and facilitate transfer to the L-TRS. This originally hypothesized similarity would fit the classical discontinuous transcription model of nidovirus sgRNA synthesis, also suggesting a conserved mechanism in *Nidovirales* to selectively enhance transcription of certain sgRNAs. Therefore, we selected three representative sgRNAs – sgRNA 2, 6 and 7 – for transcription/accumulation level analysis. Based on the aforementioned hypothesis, we expected sgRNA 7 to be the one that is upregulated most due to the existence of the RNA structure just upstream of the N ORF B-TRS. Since this RNA structure is located within the M ORF, how the RNA structure affects the transcription/accumulation level of sgRNA 6 also merited investigation. Since the B-TRS of sgRNA 2 is the one most distal from the RNA structure, we expected it to be least affected in transcription/accumulation. However, our follow-up analysis using these three representative sgRNAs revealed a very different scenario. Mutagenesis of the high-order RNA structure had an overall negative impact on sgRNA transcription/accumulation, even for the most distal sgRNA 2. Surprisingly, the relative accumulation ratio of sgRNA to gRNA was severely reduced in the stem-loop 1 mutants (5SL1 and 3SL1) and the extended stem-loop mutants (CR and extSL3), and to a lesser but nonetheless substantial extent in the Loop1 and 5SL2 mutants. In contrast, the mutant viruses only exhibited limited negative effects on full-length genomic RNA transcription/accumulation. In addition, the thermo-stability of the high-order RNA structure was shown to be important for maintaining the capability of infectious particle production. These results suggest that the integrity of the conserved RNA structure within the M ORF is critical to maintain appropriate levels of sgRNA accumulation. The phenomena we observed here obviously contradict our original hypothesis, in which the conserved RNA structure within the M ORF would function as a “road-block” to interfere with RTC processivity and promote template switching mainly for generating sgRNA 7 (5, 7).

To explain the roles of the conserved high-order RNA structure in discontinuous transcription maintenance, several hypotheses could be proposed here. (I) One possibility might be that the conserved high-order RNA structure within the M ORF regulates the stability of transcribed viral RNAs, perhaps by serving as a *cis*-acting RNA-stability element to antagonize host nonsense-mediated mRNA decay (35). The sgRNA 7 B-TRS is located just downstream of the conserved RNA structure; thus, the conserved RNA structure does not exist in sgRNA 7. Hence, the stability of sgRNA 7 at least would not be directly affected by mutations in the conserved RNA structure. However, a consistent pattern of accumulation ratio changes was observed in all three representative sgRNAs (sgRNA 2/6/7), but was not observed in the accumulation of full-length gRNA that also contains the conserved RNA structure. This argues against the conserved RNA structure mediating its effect by functioning as an RNA stability element. (II) Another possibility is that the conserved RNA structure within the M ORF functions as a regulatory RNA signal for discontinuous transcription. In the nidovirus replication cycle, there are three critical steps: 1) translation of the replicase genes (ORF1a and ORF1ab) from full-length genome; 2) synthesis of the anti-genome which is used as a template to synthesize new full-length genomes for further translation, replication and/or packaging into virions; and 3) the discontinuous synthesis of (−) sgRNAs from full-length genome, which are used as templates to transcribe (+) sgRNAs for structural protein expression and virion production (3, 4). These three steps – replicase expression, replication and transcription – all require well-coordinated and tightly controlled expression levels of gRNA and sgRNA in order to efficiently generate infectious particles. How these steps are regulated spatially and/or temporally is not fully understood. In the current study, gRNA accumulation/transcription was found to be only modestly affected by mutations in the conserved RNA structure, while the accumulation/transcription of sgRNAs was severely downregulated in a non-specific way. This suggests that nidovirus gRNA replication can be decoupled from sgRNA transcription, and the conserved RNA structure observed in the arteriviral M ORF may serve as an important regulatory signal for balancing the transcription and replication equilibrium. Whether and how host/viral trans-acting factors interact with the M ORF RNA structure to coordinate arteriviral transcription and replication requires further in-depth investigation.

We used the economically important swine virus PRRSV as a model system for our experimental investigation. Due to the very large number of sequenced isolates, PRRSV-1 and PRRSV-2 also provided the most robust comparative genomic support for functional RNA structure in the M ORF. Nonetheless, other viruses within subfamily *Variarterivirinae* (genera *Betaarterivirus*, *Nuarterivirus* and *Gammaarterivirus*; predominantly rodent and swine viruses) were also predicted to harbour the SL1, SL2, and extSL structures, with the exception of Lopma virus. Similarly, viruses in subfamily *Simarterivirinae* (simian arteriviruses) were predicted to harbour SL1 and extSL structures but not SL2 (consistent with mutations to the PRRSV-1 SL2 generally being less inhibitory than mutations to SL1 or extSL). Lopma virus has different predicted RNA structures in this region but, in the absence of comparative genomic support or an infectious clone, it is impossible to say whether or not these structures may be functional analogues of SL1, SL2 and extSL. Our current study did not extend to more divergent arteriviruses such as equine arteritis virus as preliminary inspections suggested somewhat different features in this genome region.

## ACKNOWLEDGEMENTS

We thank Eric J. Snijder (Leiden University Medical Center, Leiden, The Netherlands) for helpful discussion. This project was supported by Agriculture and Food Research Initiative competitive grant no. 2015-67015-22969 from the USDA National Institute of Food and Agriculture (to Y.F.), and grants from the Wellcome Trust (grant no. 106207 and 220814 to A.E.F).

