## Supplementary figures and images for "An intra-family conserved high-order RNA structure within the M ORF is important for arterivirus subgenomic RNA accumulation and infectious virus production"

### FigS1

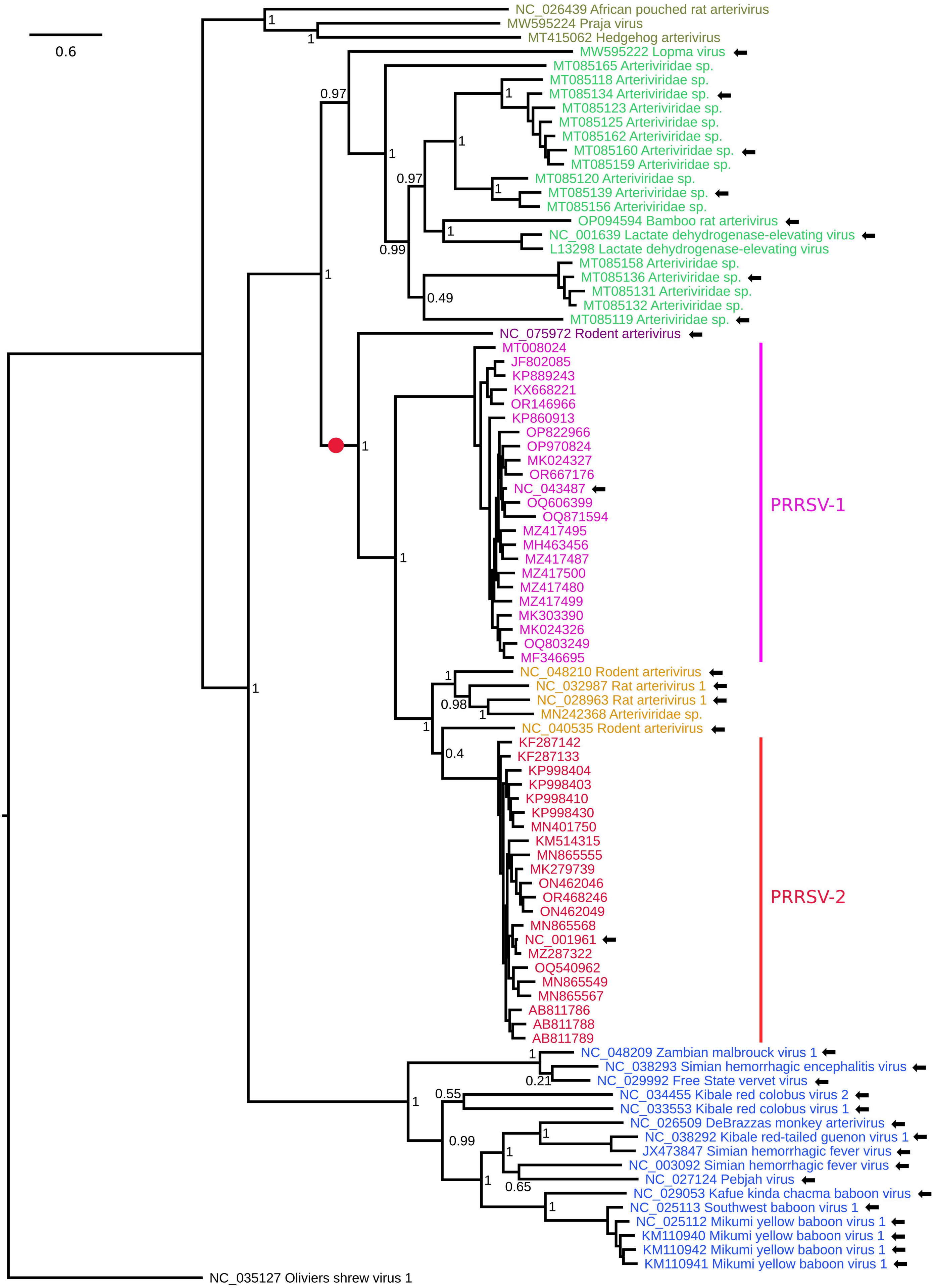
