## Supplementary material for "An intra-family conserved high-order RNA structure within the M ORF is important for arterivirus subgenomic RNA accumulation and infectious virus production": FigS2

```
MW595222        CUG CUU CGG CUA GGC CGC AGC UAC AUU UUU GCU CCU GCA AGU CAC UUA UCA ACU GAG UCA
MT085134        UUG UGC UGC UUA GGU CGC AGU UAC AUC CUA GCC CCG CCC AGU CAC GUG GAC ACC UCU GAA
MT085160        UUG UGC UGC UUA GGU CGC AAG UAC AUC CUA GCC CCG CCC AGU CAC GUG GAC ACC UCU GAA
MT085139        AUG UGC UUC UUA GGC AGA AGU UAC AUA UUA GCC CCG CCC AAC CAC GUG GAA ACA GAC GAC
OP094594        UUG UGC UGC UUA GGU CGG CGG UAC AUC UUA GCC CCG CCC AAU CAU GUU GAG AGU GUU GAU
NC_001639       UUG UGC UUC CUA GGU AGA AGU UAC AUC CUA GCC CCA CCC AGC CAC GUG GAC ACC UCU GAC
MT085136        AUG UGU UGC UUA GGU CGG CAA UAC AUC UUU GCA CCU ACC AAU CAU GUC UUA ACG GCA GAU
MT085119        AUG UGU UGC CUA GGC CGC AAA UAC AUC UUU ACC CCA AGC AAU CAC AUC GUC ACC GCA GAA
PRRSV-1         UUG UGU UGC CUU GGC CGG CGA UAC AUU CUG GCC CCU GCC CAU CAC GUA GAA AGU GCU GCA
PRRSV-2         UUG UGC UUG CUA GGC CGC AAG UAC AUU CUG GCC CCU GCC CAC CAC GUU GAA AGU GCC GCA
                 (( ((( ((( (  SL1   )) ))) ))) )            (( ((( ((  ((( ( extSL 5'

MW595222        GGC UUG CUG CCC GUG CAG --- AGC GCA AGU GCC GCC UAC GUU GUA CGG UCU CCU GGA CAG
MT085134        GGG AAA CAG AGA CUA ACC ACA UCU CAC AAC ACC GCA UUU GUG GUU AGA AAG CCA GGU UCG
MT085160        GGG AAA CAG AGG CUA ACC ACA UCU AAU AAC ACC GCA UUU GUG GUU AGA AAG CCA GGU UCG
MT085139        GGA CGU CGU GCC CUA ACC ACA UCA UCA UCC ACC GCU UUU GUG AUU AGG AGG CCG GGA UCA
OP094594        GGA CAU CAA CCA CUA ACU ACA ACU GCU GAC ACC GCA UUU GUC GUU AGA AAG CCA GGU CAC
NC_001639       GGA CGU CAG AGC CUA ACC ACA UCG UCA ACA ACC GCC UUU GUG GUU AGG AAG CCA GGU AGU
MT085136        GGA CAU CUG CCA AUA ACC ACA AAU UCU AAA UCU GCC UUU GUG GUU AGA AGG CCA GGG UCA
MT085119        GGA UGC CAC CCA AUA ACC ACU AAU UUA GAU ACU GCA UUU GUG GUU AGG AAG CCC GGC GAG
PRRSV-1         GGU CUC CAU UCA AUC UCA GCG UCU GGU AAC CGA GCA UAC GCU GUG AGA AAG CCC GGA CUA
PRRSV-2         GGC UUU CAU CCG AUU GCG GCA AAU GAU AAC CAC GCA UUU GUC GUC CGG CGU CCC GGC UCC
                                     (( ((( ((( (    SL2     )) ))) ))) )

MW595222        ACC ACC GUC AAC GGA CAA GUC GUU CCA AAG UUC AGA GCA CUG GUG CUC AAC GGC UUG AAA
MT085134        ACC CUU AUA AAC GGC CAG CUA GUG CCG AGC UUC AAG AGC CUC GUG AUU GGG GGC AGA AAA
MT085160        ACC CUU AUA AAC GGC CAG CUA GUG CCG AGC UUC AAG AGC CUC GUG AUU GGG GGC AGA AAA
MT085139        ACC ACU GUA AAC GGG ACU CUC GUC UCG GGC CUA AAA GGA CUC GUG CUU GGG GGC AGG AAA
OP094594        ACC CUU AUA AAC GGA CAA UUG GUU CCG GAC UUC AAA UCA AUU GUG AUU GGG GGC AGA AAA
NC_001639       ACC CUU GUG AAC GGG CAG UUG GUC CCG GAC UUU CAA AGA CUC GUG CUU GGG GGC AAG AAG
MT085136        ACC CUU GUA AAC GGC CAA CUG GUU CCG GAC CUG AAG AAG AUU GUG AUU GGU GGC AAA CUA
MT085119        ACU CGG GUC AAC GGU GAA UUG GUG CCC CAC AUA AAA GGG AUU GUG AUU GCU GGC AGG CGA
PRRSV-1         ACA UCA GUG AAC GGC ACU CUA GUA CCA GGA CUU CGG AGC CUC GUG CUG GGC GGC AAA CGA
PRRSV-2         ACU ACG GUC AAC GGC ACA UUG GUG CCC GGG UUG AAA AGC CUC GUG UUG GGU GGC AGA AAA
                                                             extSL 3' ) )))  )) ))) ))
```
