## Supplementary material for "An intra-family conserved high-order RNA structure within the M ORF is important for arterivirus subgenomic RNA accumulation and infectious virus production": Fig3

```
NC_048209    CGU CUG UGC CGU UUA GGA CCC CGG UAC ACG GUA GCA CCU UCG UCA UUC GUA GAG UCA ACA
NC_038293    CGU UUG UGU CGU UUA GGA CCC CGA UAC AUC ACA GCA CCU UCG UCA UUC AUU GAG UCC ACA
NC_029992    CGU CUG UGC CGC UUA GGA CCU CGG UAC ACG ACA GCA CCC UCG UCA UUC GUA GAG UCA ACA
NC_034455    CGU AUG UGU CGA CAC GGU CGG CGA UAC AUU ACG GCC CCU GCC UCU UUC AUU GAG UCC UCA
NC_033553    AGA CUA UGC CGU GUC GGC CCA CGG UAU AUG UUU AGU CCU UCC AGC UUC AUC GAG UCU UCU
NC_026509    AGA AUG UGC AGA CUC GGC CGU CAG UAC AUC ACU GCC CCU UCC AGU AUG GUC GAA ACC UCA
NC_038292    AGG AUG UGU UGG CUC GGC CGC CAA UAU GUU CUU GCC CCU UCC AGC AUG GUU GAA ACC UCA
JX473847     AGG AUG UGU UGG CUC GGC CGC CAA UAU AUU CUU GCC CCU UCC AGC AUG GUU GAA ACC UCA
NC_003092    CGG AUG UGU UGG CUC GGC CGG CAA UAC AUA ACC GCC CCU UCC AGU AUG GUU GAG UCA UCC
NC_027124    AGA AUG UGC CGU CUU GGC CGU GGC UAC AUU ACU GCC CCU UCC UCC AUG AUC GAG UCC UCC
NC_029053    AGG AUG UGU CGC CUC GGC CCG GGA UAC AUC UUA UCU UCC CCA AAC CAU GUU GAU UCC UCU
NC_025113    AGG AUG UGC CGG CUC GGC CGC CAG UAC AUC AUC UCA CCA GCA UCA CAU GUG GAA ACC UCC
NC_025112    AGG AUG UGC CGG CUC GGC CGC CAG UAC AUC AUC UCA CCA GCA UCA CAU GUG GAA ACC UCC
KM110940     AGA AUG UGC CGG CUC GGC CGC CAG UAC AUU AUC UCA CCA GCA UCA CAU GUG GAA ACC UCC
KM110942     AGG AUG UGU CGG CUC GGC CGC CAA UAC AUC GUC UCA CCA GCA UCA CAU GUG GAA ACC UCC
KM110941     AGG AUG UGU CGG CUC GGC CGC CAA UAC AUC AUC UCA CCA GCA UCA CAU GUG GAA ACC UCC
             ((( ((( ((( ((( (   SL1    )) ))) ))) ))) ))      ( ((( ((( ( extSL5'

NC_048209    UCG GGC ACG CAU ... ... UUG GUC CCU GAC GUG AAA AAG AUG GUA CUG AAU GGC AAG GUA
NC_038293    UCG GGC ACG CAU ... ... UUG GUC CCU GAC GUG AAA AAG AUG GUA UUG AAU GGC AAG GUA
NC_029992    UCA GGC ACG CAU ... ... UUG GUC CCU GAC GUG AAG AGG AUG GUA CUG AAU GGC AGG GUA
NC_034455    UUC GGU CGU CAU ... ... CUC GUC CCG GAC GUG AAG AAA AUA AUG UUA GCU GGC AGG GUU
NC_033553    UUG GGU CGC AUC ... ... CUA GUC CCG GAU GUA AAG AUG AUG GUG UUA GCU GGG AAG GUU
NC_026509    CUU GGC CGU UCG ... ... CUG GUG CCG GAU GUG AAA AGG AUC AUA CUA CAU GGA AGG GUU
NC_038292    UAC GGC CGU UCG ... ... CUG GUG CCG GAU GUG AAA CGU AUA AUC UUG AAU GGA AGG GUU
JX473847     UAC GGC CGU UCG ... ... CUG GUG CCG GAU GUG AAA CGU AUA AUC UUG CAU GGA AGG GUU
NC_003092    CUU GGC CGU UUA ... ... CUC AUG CCG GAU GUG AAA AGG AUC AUA CUC AAU GGA AGG GUU
NC_027124    GCA GGC CAU CAC ... ... UUG GUU CCG GAC GUU AAG CGG AUC AUA CUC CAA GGA AGG GUU
NC_029053    CUU GGC CGU UAU ... ... CUC AUC CCC GAC GUG AAA AGA AUG GUU CUA GCA GGG AAG AUU
NC_025113    UUU GGC CGU UAU ... ... CUC AUC CCG GAC GUG AAA AAG UUA GUG CUU GCU GGU AAG AUU
NC_025112    UUU GGC CGU UAU ... ... CUC AUC CCG GAC GUG AAA AAG UUA GUG CUU GCU GGU AAG AUU
KM110940     UUU GGC CGU UAU ... ... CUC AUC CCG GAC GUG AAA AAG UUA GUG CUU GCU GGU AAG AUU
KM110942     UUU GGC CGU UAU ... ... CUC AUC CCG GAC GUG AAA AAG UUA GUG CUU GCU GGU AAG AUU
KM110941     UUU GGC CGU UAU ... ... CUC AUC CCG GAC GUG AAA AAG UUA GUG CUU GCU GGU AAG AUU
                                                                 extSL 3' ) ))) ))) )
```
