## Supplemental_figure_legends for "An intra-family conserved high-order RNA structure within the M ORF is important for arterivirus subgenomic RNA accumulation and infectious virus production"

**Figure Legends for Supplemental Figures**

**Figure S1. Arterivirus phylogenetic tree.** The tree is based on the ORF1ab amino acid sequences from 91 representative arterivirus sequences and was produced with PhyML (see Methods). The divergent wobbly possum disease virus and equine arteritis virus clades were excluded. Values at nodes indicate approximate likelihood supports as estimated by PhyML. Sequences used in Figures 4A, S2 and S3 are marked with black arrows. The red dot marks the *Betaarterivirus* (PRRSV-1, PRRSV-2, some rodent arteriviruses) and *Nuarterivirus* (NC_075972) clade. Note that the divergent sequence MT085165 was not included in Figure S2 because it is an incomplete sequence and lacks the M and N ORF regions.

**Figure S2. Potential conservation of SL1, SL2 and extSL in lactate dehydrogenase-elevating virus (genus *Gammaarterivirus*) and relatives, but not in Lopma virus (MW595222).** The SL1-SL2-extSL region of selected sequences in the clade colored in green in Figure S1 was extracted and aligned to the corresponding region of PRRSV-1 and PRRSV-2 (blue text). The base pairings predicted in NC_001961 (PRRSV-2) are highlighted in yellow (SL1), orange (SL2) or green (extSL), except that G:U base pairs are indicated by highlighting either the G or the U in cyan. Compensatory substitutions in NC_043487 (PRRSV-1) are highlighted in pink. Potential structures corresponding to SL1, SL2 and extSL in the other sequences are annotated using a similar color scheme, but additional unrelated base pairings (e.g. offset via insertions or deletions) are annotated in shades of blue and brown, and additional compensatory substitutions are annotated in dark orange, purple and red. Additional potential structure in other regions of the alignment is not shown. M ORF codons are separated by spaces.

**Figure S3. Potential conservation of SL1 and extSL in subfamily *Simarterivirinae*.** The SL1-extSL region of all representative sequences in the clade colored in blue in Figure S1 was extracted from an M ORF alignment. M ORF codons are separated by spaces. For SL1, the base pairings conserved across the alignment ("core SL1") are highlighted in brown whereas light pink denotes flanking base pairings that are supported by RNAfold but offset relative to the "core SL1" due to unpaired bulge nucleotides. G:U base pairs are indicated by highlighting either the G or the U in cyan. Compensatory substitutions in the "core SL1" are highlighted in pink or green. For extSL, several distinct (i.e. non-aligning) structures are predicted by RNAfold in different lineages, colored mauve, slate blue, yellow and green. Compensatory substitutions within a structure group are highlighted in pink, and G:U base pairs are indicated by highlighting either the G or the U in cyan. Some additional unaligned potential base pairings are highlighted in olive green. Additional potential structure in other regions of the alignment is not shown. The "... ..." represents the excision of a 23-codon alignment block (22 or 23 codons depending on the sequence) to reduce the size of the figure.
